# Foundation Models Meet Imbalanced Single-Cell Data When Learning Cell Type Annotations

**DOI:** 10.1101/2023.10.24.563625

**Authors:** Abdel Rahman Alsabbagh, Alberto Maillo Ruiz de Infante, David Gomez-Cabrero, Narsis A. Kiani, Sumeer Ahmad Khan, Jesper N. Tegnér

## Abstract

With the emergence of single-cell foundation models, an important question arises: how do these models perform when trained on datasets having an imbalance in cell type distribution due to rare cell types or biased sampling? We benchmark three foundation models, scGPT, scBERT, and Geneformer, using skewed single-cell cell-type distribution for cell-type annotation. While all models had reduced performance when challenged with rare cell types, scGPT and scBERT, performed better than Geneformer. Notably, in contrast to scGPT and scBERT, Geneformer uses ordinal positions of the tokenized genes rather than actual raw gene expression values. To mitigate the effect of a skewed distribution, we find that random oversampling, but not random undersampling, improved the performance for all three foundation models. Finally, scGPT, using FlashAttention, has the fastest computational speed, whereas scBERT is more memory-efficient. We conclude that tokenization and data representation are essential areas of research, and new strategies are needed to mitigate the effects of imbalanced learning in single-cell foundation models. Code and data for reproducibility are available at https://github.com/SabbaghCodes/ImbalancedLearningForSingleCellFoundationModels.

## 1 Introduction

The advent of high-performance computing has facilitated the development and implementation of advanced Large Language Models (LLMs) such as GPT-4 [1] and LLaMA [2], marking significant progress in various fields. These LLMs exhibit a unique ability to utilize transfer learning, capitalizing on knowledge acquired from generic domains as a base to build understanding in specific domains, showcasing superior performance compared to models trained solely on domain-specific data. The exploration and successful incorporation of these LLMs have set the stage for unprecedented advancements in diverse scientific domains. The notion of foundational models may constitute the next logical step beyond tailor-made task-specific bioinformatics and modeling pipelines [3, 4].

In single-cell biology, the introduction of single-cell foundation models has marked a significant advancement in deciphering cellular diversity [5]. These models, such as scGPT [6], scBERT [7], and Geneformer [8], have displayed versatility in analyzing and interpreting the single-cell data, potentially overcoming the limitations inherent to earlier, task-specific model architectures. Yet, it remains unclear how to evaluate and validate the performance of such foundational models beyond specific downstream tasks. Here, we ask how resilient such models are to skewed, imbalanced data distributions. This problem is, in our hands, a generic challenge in the biomedical domain due to selective samples and the occurrence of rare data types.

This paper presents a focused comparative analysis of three recent single-cell foundation models, exploring their efficacy and limitations in handling imbalanced learning targeting cell-type annotation. Our experiments are conducted using the Multiple Sclerosis (MS) [9] and Zheng68k [10] datasets (Section 2.2), two well-known datasets for assessing cell type annotation performance. We also examine the effect of various data-driven sampling techniques, such as random undersampling, oversampling, and imputation (Section 2.3), to address and potentially mitigate the effects of such imbalances. We analyze the correlation between modified data distribution and the overall performance of the models. Our results show that the performance of these models is indeed affected when applied to imbalanced datasets and that oversampling enhances their predictive performance for cell-type annotation tasks by equalizing the cell-type distribution.

## 2 Methods

This section explains the methods used throughout this study and the applied sampling technique to solve the data imbalance issue in detail.

### 2.1 Foundation Models

#### scGPT

scGPT is a dedicated variant of the GPT model, explicitly customized for single-cell data with a specially designed attention mask. It undergoes pretraining using an extensive dataset of more than 10M cells sourced from public scRNA-seq data spanning six different tissues and has approximately 50M trainable parameters. Cross entropy loss (equation 1) was utilized as a loss function.

#### scBERT

scBERT is a specialized Large Language Model (LLM) built on BERT and tailored for single-cell genomics. Built upon the Performer architecture and initialized with gene embeddings from Gene2vec, it’s pretrained on over 1M cells from public scRNA-seq data from 74 tissues and with around 10M learnable parameters. The cross-entropy loss (equation 1) function was employed.

#### Geneformer

Geneformer is an LLM designed to process ranked gene expression values in single cells, treating them as tokenized strings where gene rankings serve as individual tokens. It goes through a pretraining phase using a vast dataset comprising over 30 million cells obtained from publicly available data spanning 39 distinct organs (specific tissue coverage is not reported). The model encompasses approximately 30M trainable parameters. Cross entropy loss (equation 1) was used as the loss function.

The cross entropy *L*(*y, ŷ*) is defined as follows:

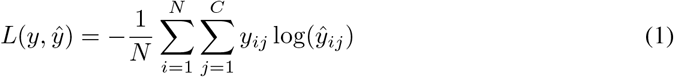

Where N is the number of samples, C is the number of classes, *y*_*ij*_ is the true label for the *i*th sample and *j*th class (0 or 1). *ŷ*_*ij*_ is the predicted probability that the *i*th sample belongs to the *j*th class. In this case, the classes are the cell types, which will be called interchangeably.

### 2.2 Datasets

#### Multiple Sclerosis (MS)

We utilized scGPT’s preprocessed version, featuring 7,844 training cells and 13,468 test cells, using the top 3,000 highly variable genes across 19 cell types. This dataset suits our study due to its class imbalance, as seen in Figure 1 (left).

**Figure 1:**
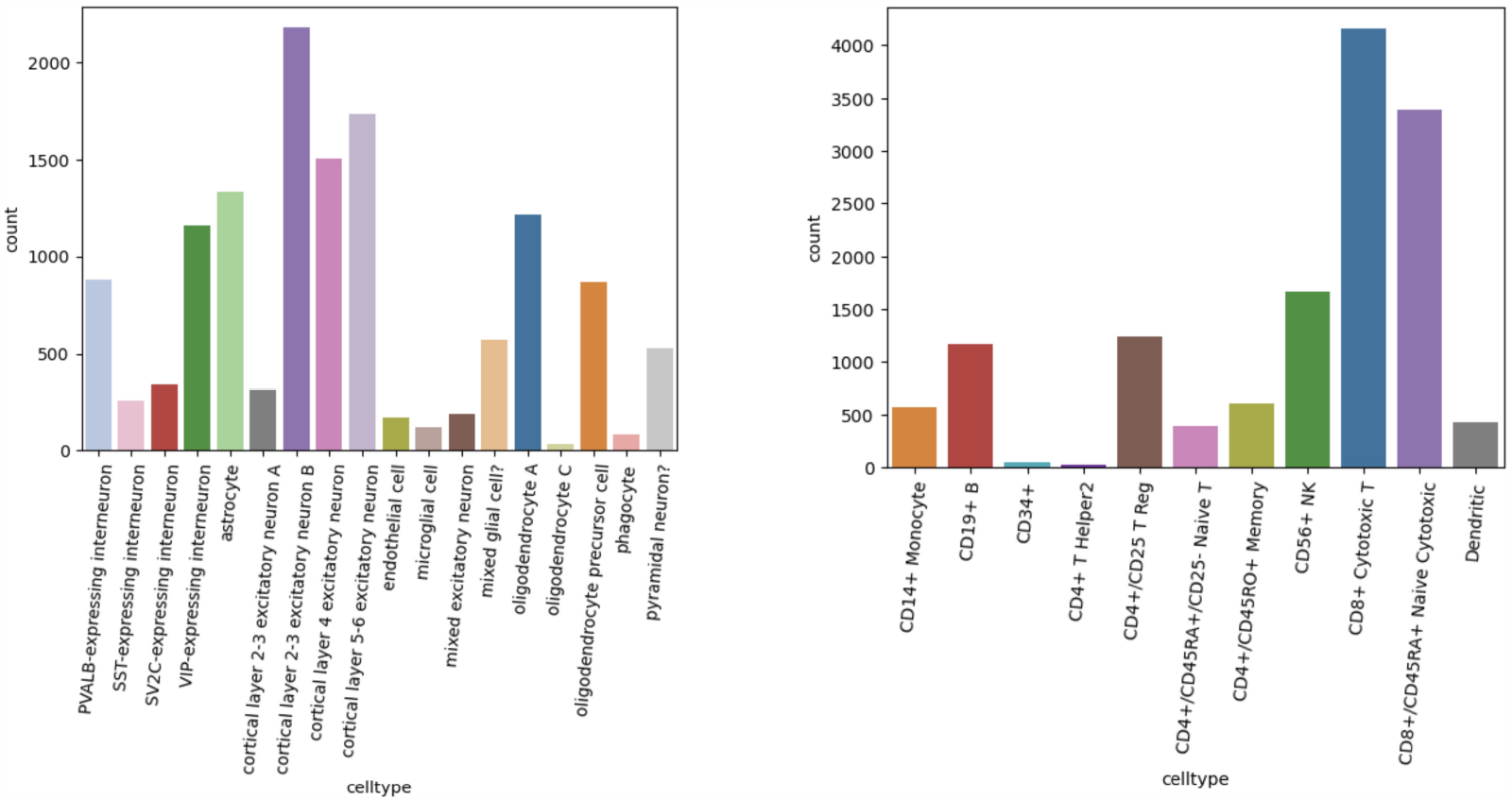
Cell type distribution of the train sets: MS dataset (left) and Zheng68K (right).

#### Zheng68K

A subset of the human PBMC, significantly larger than MS, comprising 54,760 training cells and 13,690 test cells, with 16,906 genes across 11 distinct cell types. Its class imbalance, as shown in Figure 1 (right), renders it a good choice for testing alongside the MS dataset.

### 2.3 Sampling Techniques

#### Random Undersampling

We performed random undersampling on the MS and Zheng68K training sets using a threshold of 300. Cell types with counts exceeding this threshold were downsized to meet it, while those below remained unchanged. This led to training set sizes of 2,990 and 2,975 instances for MS and Zheng68K, respectively.

#### Random Oversampling

We conducted random oversampling on both datasets by setting the oversampling threshold to the count of the majority cell type—2,010 for MS and 16,600 for Zheng68K. Consequently, the training sets expanded to 38,190 and 182,600 instances, respectively.

#### Oversampling With Imputation

We oversampled with imputation using scVI [11] for both datasets, setting thresholds at the average count of the main cell types: 1,090 for MS and 6,053 for Zheng68K. We increased the count of cell types below these thresholds until they met, resulting in training sizes 20,734 for MS and 85,381 for Zheng68K.

## 3 Experimental Setup

We used default hyperparameters and model settings from each paper, where scGPT and scBERT utilize Adam with a learning rate of 10^−4^, *β*_1_ and *β*_2_ of 0.9 and 0.999 respectively, whereas Geneformer used AdamW as the optimizer with a learning rate of 5 *×* 10^−5^ and the same *β*_1_ and *β*_2_ as the former models. All three models used learning rate schedulers, specifically step, cosine annealing, and linear schedulers for scGPT, scBERT, and Geneformer, correspondingly, and with the same order, the batch sizes were set to 32, 1, and 12. scBERT used a stratified shuffle split with 5 splits and a validation split size of 20% (out of the train set), the split with the lowest validation loss was chosen. Similarly, scGPT has a validation split size of 10%. We kept the objective of minimizing the cross entropy loss the same as in all papers. The gene length hyperparameter was changed for each dataset, particularly 3,001 for the MS dataset, and 16,907 for the Zheng68K dataset, with an increase of 1 of the original gene number to specify the class token. Experiments were run on an NVIDIA A100 GPU with 64GB of memory, with Python version 3.8.17 and CUDA version 11.7. We find that these software versions universally work on all three models. One solution to the variations of dependencies across the models was to use the 22.08-py3 PyTorch Docker container, which helped significantly, especially with importing the transformer model called FlashAttention [12] that takes advantage of complex parallelism settings underneath, which makes the learning operations faster and more memory-efficient. We assessed the performance of the models using the following metrics: accuracy, precision, recall, and macro F1-score.

## 4 Results & Discussion

We evaluated the performance of three open-source single-cell foundation models scGPT, scBERT, and Geneformer. We assessed their performance on cell-type annotation tasks on MS and Zheng68K datasets. We discuss the effect of class imbalance in terms of cell types on the performance of these single-cell foundation models. We also examined the effectiveness of various sampling techniques in addressing these issues on model performance.

### scGPT and scBERT exhibit a comparable level of performance on the default datasets, whereas Geneformer notably lags behind

Figure 2, with exact values enumerated in Table 1 (also see Figure 4) shows that scGPT and scBERT perform comparably well (F1-score of 0.753 and 0.858) on the MS dataset respectively, while Geneformer trails with 0.560. This trend is consistent in the Zheng68K dataset, with scGPT’s and scBERT’s F1-score of 0.725 and 0.743, and Geneformer 0.684. We posit that the observed discrepancies result from Geneformer’s unique input preprocessing methodology. In this model, genes are represented as tokens but ordered in descending order based on their gene expression values, focusing solely on ordinal positions. In contrast to scGPT and scBERT, Geneformer ignores the actual gene expression values. Additionally, we visualized the predictions using the UMAP plots (Figure 3) for the qualitative evaluation of these foundation models. The UMAP plots show that the rare cell types suffer from misclassification specifically Oligodendrocyte C and Phagocyte in MS and CD34+, CD4+ T Helper2 in Zheng68K, thereby emphasizing the imbalance learning issue with these models. This issue is more clearly depicted in Figure 5, where the confusion matrices reveal the cell types that are distinctly misclassified in all three models, which are the minority classes as shown in the dataset distributions in Figure 1. In summary, scBERT generally does slightly better than scGPT, especially when no changes are made to the datasets, showing it is better at dealing with imbalanced learning problems. In contrast, Geneformer is more likely to overfit than scBERT, as seen in Figure 6 (a), where scBERT’s loss steadily and monotonically decreases, unlike Geneformer. scGPT also overfits, but not as severely as Geneformer does. On an overall level, we notice that scBERT is the favorable choice for a foundation model when dealing with imbalanced data.

**Table 1:** Results after running the foundation models on different sampling techniques on MS and Zheng68K datasets, with bold text referencing the best intra-model performance on the sampling techniques and green highlight accentuating the best overall inter-model performance.

| Dataset |  | Multiple Sclerosis (MS) |  |  |  | Zheng68K |  |  |  |
| --- | --- | --- | --- | --- | --- | --- | --- | --- | --- |
| Model | Setting | Accuracy | Precision | Recall | Macro F1 | Accuracy | Precision | Recall | Macro F1 |
| scGPT | Default | 0.864 | 0.811 | 0.764 | 0.753 | 0.840 | 0.733 | 0.725 | 0.725 |
|  | Random Undersampling | 0.829 | 0.779 | 0.836 | 0.786 | 0.666 | 0.589 | 0.714 | 0.616 |
|  | Random Oversampling | **0.964** | **0.954** | **0.970** | **0.962** | **0.964** | **0.954** | **0.985** | **0.968** |
|  | Oversampling With Imputation | 0.853 | 0.819 | 0.759 | 0.760 | 0.847 | 0.739 | 0.767 | 0.749 |
| scBERT | Default | 0.872 | 0.910 | 0.850 | 0.858 | 0.769 | 0.753 | 0.739 | 0.743 |
|  | Random Undersampling | 0.859 | 0.851 | 0.811 | 0.818 | 0.673 | 0.671 | 0.643 | 0.642 |
|  | Random Oversampling | **0.965** | **0.966** | **0.965** | **0.965** | **0.960** | **0.960** | **0.960** | **0.959** |
|  | Oversampling With Imputation | 0.929 | 0.936 | 0.929 | 0.931 | 0.757 | 0.771 | 0.749 | 0.757 |
| Geneformer | Default | 0.731 | 0.550 | 0.587 | 0.560 | 0.783 | 0.742 | 0.661 | 0.684 |
|  | Random Undersampling | 0.618 | 0.483 | 0.517 | 0.494 | 0.500 | 0.509 | 0.466 | 0.454 |
|  | Random Oversampling | **0.764** | **0.646** | **0.627** | **0.620** | **0.957** | **0.957** | **0.956** | **0.955** |
|  | Oversampling With Imputation | 0.666 | 0.517 | 0.558 | 0.525 | 0.777 | 0.692 | 0.647 | 0.659 |

**Figure 2:**
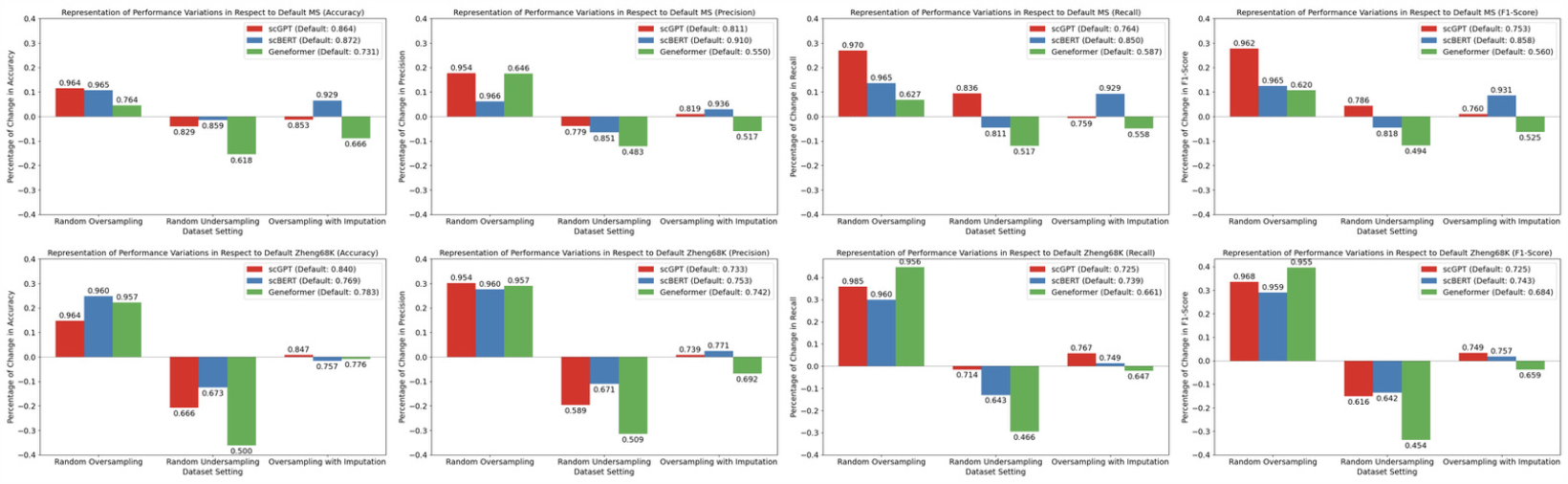
Representation of performance variations, expressed in percentage, with different sampling techniques compared to default setting. Datasets are presented as rows with MS (first) and Zheng68K (second), and evaluation metrics as columns with accuracy (first), precision (second), recall (third), and F1-score (fourth). Default results in the legend for each sub-figure.

**Figure 3:**
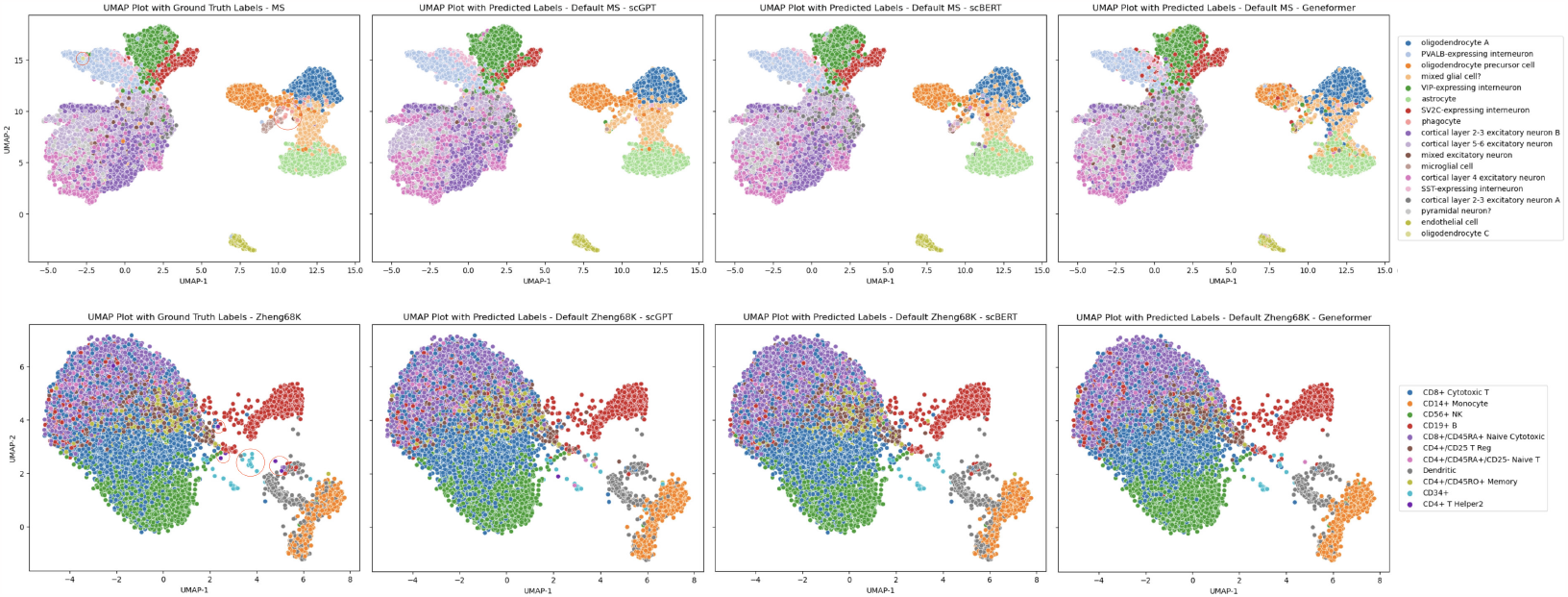
UMAP plots representing the MS (first row) and Zheng68K (second row) test sets following the application of foundation models on the default datasets. The columns display the ground truths (first), along with the predictions made by scGPT (second), scBERT (third), and Geneformer (fourth). In the ground truth plots, rare cell types are distinctly encircled in red.

**Figure 4:**
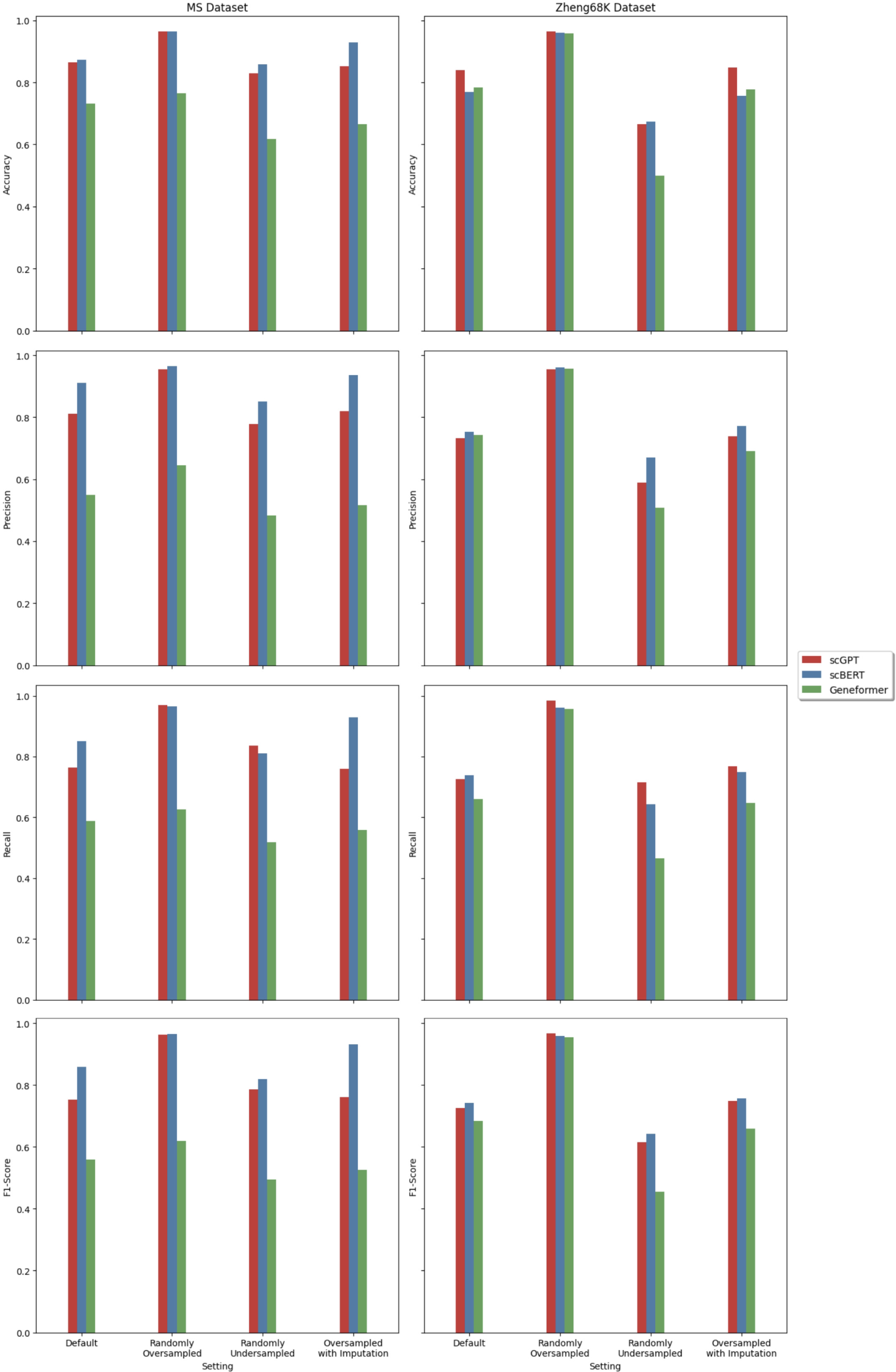
Results after running the foundation models on different sampling techniques, with datasets as columns with MS (left) and Zheng68K (right), and evaluation metrics as rows with accuracy (first), precision (second), recall (third), and F1-score (fourth).

**Figure 5:**
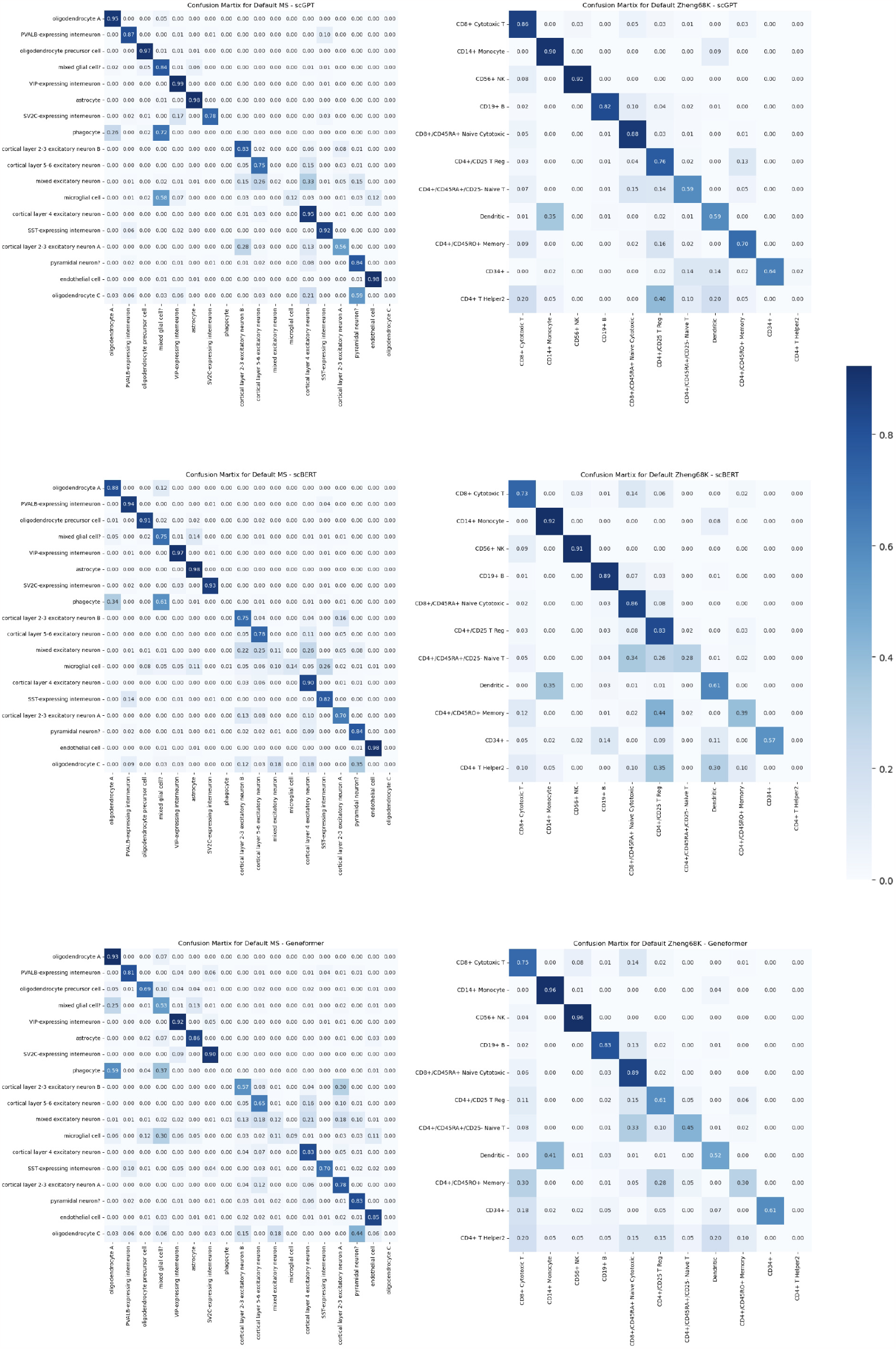
Confusion matrices after running the foundation models on MS (first column) and Zheng68K (second column) test sets, with each row having a different foundation model: scGPT (first), scBERT (second), and Geneformer (third).

**Figure 6:**
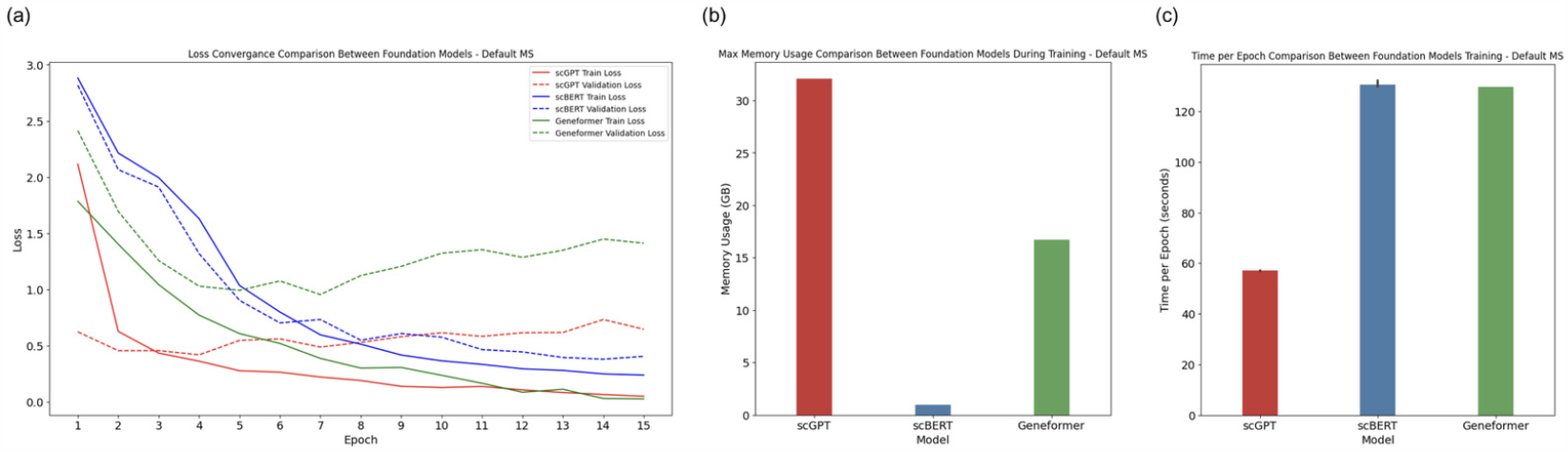
Comparative performance of scGPT, scBERT and Geneformer on the default MS dataset: showing (a) loss convergence of training and validation losses, (b) time taken per training epoch, and (c) maximum memory usage while training.

### Random oversampling consistently yielded the best performance across all foundation models

Random oversampling of the minority classes in the MS and Zheng68K datasets significantly enhanced all models’ performance. As highlighted in Figure 2 (see full illustration in Figure 4), oversampling stands out as an effective strategy for overcoming challenges posed by imbalanced learning. Oversampling achieved an accuracy of ∼96% on almost all cases, showing an improvement of around ∼5%-∼10% on MS and ∼15%-∼25% on Zheng68K in accuracy, compared to the results obtained from the original datasets. Moreover, compared to the original datasets, we noted increments of approximately ∼ 10%- ∼ 30% in F1-score on the MS, and ∼ 30%- ∼ 40% on Zheng68K dataset. Interestingly, scGPT performs better with the oversampled MS dataset. In contrast, Genformer performs better than scGPT and scBERT on the oversampled Zheng68k dataset, which substantiates that the poor performance of Geneformer can be mitigated with a more extensive and balanced dataset. In the second setting, undersampling the majority classes, the foundational models exhibited poor performance with a decrease in accuracy ranging from ∼ 2%- ∼ 20% on MS, and ∼ 13%- ∼ 38% on Zheng68K, with Geneformer being the most affected in both experiments. Additionally, oversampling with imputation slightly impacted scGPT but improved scBERT significantly while adversely affecting Geneformer. Both random undersampling and oversampling with imputation performed similarly on the MS dataset, likely due to its smaller size and variability than Zheng68K. The subpar performance of oversampling with imputation may stem from their inability to accurately represent the actual data distribution, whereas oversampling directly addresses imbalances, offering a more favorable learning environment and improved performance. Even though random oversampling is just a process of replication of the minority classes, it plays an important role in mitigating the natural bias towards the majority class issue, which appears - from results observation - that all foundation models implicitly suffer from and can’t address properly. In contrast, although random undersampling improves class balance similar to random oversampling, it sacrifices important information held within the majority classes, causing a noticeable performance degradation.

### scGPT exhibits the fastest computational speed, albeit at the cost of being the most memory-intensive

In Figure 6 (b) and (c), it is shown that scGPT works about twice as fast as both scBERT and Geneformer, requiring only about a minute for each epoch as it uses the faster FlashAttention method. However, scBERT uses much less memory, about 30 times less than scGPT and 16 times less than Geneformer. Thus, for memory efficiency, scBERT is ideal; for time efficiency, scGPT is preferable.

## 5 Conclusion

With the emergence of foundational models in genomics come new challenges. For example, assessing and benchmarking their performance on different downstream tasks is critical. This is a stepping stone towards delineating relevant metrics for evaluating such foundation models. To this end, we have assessed three recent foundation models by challenging them with skewed data distributions. The difference in the data representation, ranked ordinal (Geneformer) versus using the expression values (scBERT and scGPT) is most likely a key element in explaining the performance difference in the context of imbalanced data distributions. This suggests how to design the initial tokenization and data representation in foundation models is an essential area for future research. As a first step towards mitigating the effects of skewed data, we find that random oversampling significantly enhances all foundation models’ performance for cell-type annotation tasks. We suggest exploring strategies, such as algorithmic approaches, including loss function engineering, or data-driven techniques, including different sampling methods to mitigate the effects of imbalanced learning in single-cell foundation models. These avenues of investigation hold promise for enhancing foundation model performance amidst data class imbalances. Beyond addressing specific downstream tasks, we anticipate that the concept of foundational models may open the path toward constructing data-driven systems biology models by leveraging learned efficient representations [13].

